# Training neural networks from scratch in a videogame leads to brittle brain encoding

**DOI:** 10.1101/2025.11.28.691119

**Authors:** François Paugam, Basile Pinsard, Marie St-Laurent, Guillaume Lajoie, Lune Bellec

## Abstract

Recent brain-encoding studies using videogame tasks suggest that the training objective of an artificial neural network plays a central role in how well the network’s representations align with brain activity. This study investigates the alignment of artificial neural network activations with brain activity elicited by a video game task using models trained from scratch in controlled settings. We specifically compared three model training objectives: reinforcement learning, imitation learning, and a vision task, while accounting for other potential factors which may impact performance such as training data and model architecture. We tested models on brain encoding, i.e. their ability to predict functional magnetic resonance imaging (fMRI) signals acquired while human subjects played different levels of the video game Super Mario Bros. When tested on new playthroughs from the game levels seen at training, the reinforcement learning objective had a small but significant advantage in brain encoding, followed by the imitation learning and vision models. We hypothesized that brain-aligned representations would emerge only in task-competent models, and that the specific brain regions well encoded by a model would depend on the nature of the task it was trained on. While brain encoding did improve during model training, even an untrained model with matching architecture approached the performance of the best models. Contrary to our hypotheses, no model layers or specific training objectives aligned preferentially with specific brain areas. Large performance gaps also persisted in fully trained models across game levels, both those seen during training and entirely novel ones. Overall, even though reinforcement learning presented a small advantage to train brain encoding models for videogame data, all tested brain encoding models exhibited brittle performance with limited generalization both within- and out-of-distribution. Overall, our results suggest that training small artificial models from scratch is not sufficiently reliable, and that incorporating pretrained models such as foundation vision–action models may ultimately be necessary to support robust inferences about brain representations.

## Introduction

Brain encoding has become a widely used technique for modeling brain dynamics during naturalistic tasks (Yamins et al. 2014; Seeliger et al. 2021). By leveraging computational models trained on rich sensory datasets, this approach provides new opportunities to study complex stimuli and their intricate relationships with neural activity. For instance, activations from models trained to identify objects from pixel-level image data can successfully predict neural responses in subjects exposed to the same images (Schrimpf et al. 2018). Brain encoding studies have demonstrated functional similarities between artificial neural networks (ANNs) and the human brain across multiple domains, including vision (Yamins and DiCarlo 2016; Cichy et al. 2016; Güçlü and Gerven 2015), auditory processing (Kell et al. 2018; Freteault et al. 2023), and language comprehension (Caucheteux and King 2022; Schrimpf et al. 2020). However, most models used for brain encoding rely on passive perceptive tasks, such as participants viewing images (Allen et al. 2022) or reading a text (Toneva and Wehbe 2019).

Expanding brain encoding to active tasks involving interaction with virtual or embodied environments marks a crucial step toward modelling complex cognition (Paolo et al. 2024; Zador et al. 2023). Video games, in particular, offer a rich framework for studying these dynamics, as they simultaneously engage visual perception, decision-making, and motor output in a highly interactive context (Burelli and Dixen 2024; Bellec and Boyle 2019) and can be used inside a magnetic resonance imaging (MRI) scanner (Harel et al. 2023). The choice of training tasks for ANNs has a significant impact on the representations the models learn. Many different types of training tasks can be applied to the stream of video game data, but it remains unclear which approach yields models that best align with brain activity, either as a whole or for specific brain networks.

Purely perceptive visual ANNs that do not predict videogame actions are expected to exhibit fewer similarities with brain regions beyond the visual cortex, compared to action models trained to optimize gameplay from the game frames. This hypothesis is supported by findings from Cross and colleagues (Cross et al. 2021), who compared a basic vision model trained for auto-encoding with a model trained through reinforcement learning (RL) in Atari games (Space Invaders, Enduro, Pong). In contrast, we observed in a previous study that a vision model pretrained on Imagenet for classification outperformed an RL-trained model optimized for gameplay in brain encoding across all brain regions in the game Super Mario Bros (Paugam et al. 2024). Finally, (Kemtur et al. 2023) used imitation learning to train models to replicate human players’ actions in the game Shinobi III. They found that models trained on an individual subject’s gameplay data achieved better brain encoding accuracy for that subject compared to models trained on another subject’s gameplay, or pure vision layers of their model which do not account for dynamics of the game. Additionally, they reported a hierarchical functional correspondence between model layers and brain regions, a result in line with previous work with static images stimuli, focused on the ventral visual system (Eickenberg et al. 2017; Yamins and DiCarlo 2016).

The apparent inconsistencies between these studies are difficult to interpret, as each study focused on different types of models and relied on different games, making direct comparisons challenging. The comparison of models based on brain encoding may also reflect other factors than the tasks used to train ANNs. For instance, in (Paugam et al. 2024) we compared models that varied not only in training objectives but also in the nature and size of their training data. For vision models, the diversity and size of the training data can have a large impact both on image annotation performance and brain encoding accuracy (Conwell et al. 2023), and a similar result was found specifically for video transformer models trained for videogame data (Ahmadi et al. 2024). The difference observed between the vision model and the RL model in (Paugam et al. 2024) may thus reflect that the vision model was trained on a very large and diverse dataset of natural images (ImageNet) while the RL model was trained on a comparatively smaller number of videogame frames all drawn from Super Mario Bros. Another potential confounding factor is evident in both (Cross et al. 2021) and (Kemtur et al. 2023), as both studies did not compare equivalent layers for encoding between vision and action models. In vision, the choice of layer in artificial neural networks impacts which brain regions it encodes optimally, regardless of model architecture (Conwell et al. 2023). So the reported differences between vision and action models may reflect the choice of layers used in the model comparison rather than the task used to train the networks.

In summary, even though previous studies have explored various training objectives for ANNs in videogame-based brain encoding, their experiments entangled multiple factors, such as model architectures or training datasets, limiting the conclusions that can be drawn on the specific impact of the training objective. With the current work, our main objective was to explore how the choice of training objective impacts an ANN’s ability to perform brain encoding of functional magnetic resonance imaging (fMRI) data, while carefully controlling for the model architecture and the data distributions used to train each model.

To do so, we compared models with matching convolutional neural network (CNN) architectures trained from scratch with the same input data distribution: frames from the super mario bros game, and only changed the training objectives between models. The different training tasks we used are as follows:

- **Proximal Policy Optimisation (PPO)**: training the model to play the game by maximizing a reward with RL through the PPO algorithm (Schulman et al. 2017),
- **Imitation learning:** the model predicts human actions from visual features using gameplay specific to a given individual,
- **Resnet proxy:** learning to predict latent features from visual features, where latent features were extracted from one layer of a pretrained image classification model called ResNet152v2 (He et al. 2015), which is known to have good performance for brain encoding (Paugam et al. 2024). This training task created a proxy of that pretrained model purely trained on our videogame dataset.

We also included two baselines: an Untrained model with similar architecture, and a **pretrained** vision model (ResNet152v2) trained on Imagenet (Deng et al. 2009).

Our first aim was to **identify which training objectives resulted in better brain activity predictions across different regions**. Our hypothesis was that *action models (RL and imitation) would outperform pure vision and baseline models, in particular in sensorimotor and attentional brain networks*.

Our second aim was to **test for the existence of parallel hierarchies of representations between models and the brain**. *Our hypothesis was that early layers would map to visual brain regions, while late layers would encode better associative and sensorimotor cortices*.

Our third aim was to **examine whether the performance of ANNs trained to generate actions was associated with their ability to model the brain**. In vision, performance on image annotation tasks correlates strongly with brain alignment, up to a certain level of accuracy (Yamins et al. 2014; Schrimpf et al. 2018). We thus hypothesized that *both the ability to play the game and the quality of behavioural imitation would be predictive of the quality of brain encoding*. This relationship should hold whether comparing the same model at different stages of training (and thus with different task performance), as well as when comparing a trained model across different levels (with, again, different task performance).

Our fourth and last objective was to **evaluate how brain encoding models generalized on out-of-distribution data**. Classical models trained through RL are notoriously brittle, and fail to generalize their behaviour to new game levels (Jiang et al. 2022; Nichol et al. 2018). We hypothesized that *the performance of brain encoding with the RL model would be severely decreased when evaluated on new game levels unseen during training, while other networks which capture more abstract and robust features, notably the ResNet152v2 model, would be more robust to such out-of-distribution evaluation*.

## Methods

### Dataset

#### Participants

This study utilizes the *Mario* dataset, publicly available through the Courtois NeuroMod (CNeuroMod) project. As not all participants in the broader project completed every task, we focused on those who completed the *Mario* protocol. This dataset explores behavioral and neural correlates of video game play and includes five participants (sub-01, sub-02, sub-03, sub-05, and sub-06; two females, three males) who played *Super Mario Bros.* (*SMB*; Nintendo, 1985) inside an MRI scanner.

While not a central focus of this study, prior videogame experience may have influenced the results. Two participants (sub-01 and sub-02) reported regular videogame play and one participant (sub-06) reported no regular videogame practice but played another retro platformer called Shinobi III: Return of the Ninja Master (RotNM; Sega, 1993) that she practiced for 10+hours as part of a prior CNeuroMod experiment (shinobi dataset) (Boyle et al. 2020).

#### MRI acquisitions and processing

About 15 hours of fMRI data were acquired per participant on a 3T Siemens Prisma Fit scanner (temporal resolution: TR = 1.49 s; spatial resolution: 2 mm isotropic) and preprocessed with *fMRIprep* (Esteban et al., 2019; version v20.2 LTS). After spatial smoothing (FWHM = 5 mm), functional MRI time series were denoised using Nilearn with the following strategy:

...

load_confounds(nifti_path, strategy=[“motion”, “high_pass”, “wm_csf”, “global_signal”], motion=“basic”, wm_csf=“basic”, global_signal=“basic”, demean=True)

...

Functional MRI data were then projected onto the MIST atlas using the *NiftiLabelsMasker* in Nilearn. We selected the MIST atlas variant with the highest resolution (1097 parcels), covering the grey matter, including the cerebellum, basal ganglia, and thalamus (Urchs et al., 2019).

#### Videogame task

The participants were recorded while they played 22 levels of *SMB,* a side-scrolling platformer developed by Nintendo. The general objective of the game is to progress rightward through each level while avoiding obstacles and enemies. In the initial *discovery* phase, participants played each level in the original game sequence, with unlimited attempts until they completed it once. Once all 22 levels were unlocked, participants entered a *practice* phase, in which levels were presented in a random order. For each level, participants had a single attempt with up to three lives to complete it before a new level was randomly selected.

#### Videogame set-up

All participants used a custom MRI-compatible video game controller, designed by the CNeuroMod team using 3D-printed plastic and fiber optics. The controller connected via USB to the stimulation computer (Harel et al. 2023). The video game was played on a console emulator using OpenAI’s *gym-retro* library (Nichol et al. 2018), a Python-based platform supporting emulators for over 10 retro consoles and thousands of games. Built on *gym* (Brockman et al. 2016), a reinforcement learning library, *gym-retro* integrates console emulators via the Libretro API (https://www.libretro.com/). This setup enables both artificial and human agents to play retro games from saved states while providing Python API access to the game’s RAM. *SMB* was played and recorded at 60 Hz. Since the game is fully deterministic, only player inputs (button presses) were recorded, allowing for precise reconstruction of gameplay video streams.

#### Train/validation/test splits

fMRI data, button presses, and frame data from 20 of the 22 levels were divided into training, validation, and test sets. The training, validation, and test sets consisted of 80%, 10%, and 10% of the game runs, respectively, maintaining similar proportions of runs from each of the 20 levels and a balanced distribution of completed/failed runs. The remaining two held-out levels were used as an out-of-distribution set to assess model generalizability.

### Model Inputs

For all models except ResNet, the input is a sequence of 16 consecutive frames (60 Hz), preprocessed using the mapping function introduced by (Mnih et al. 2015), a widely used method for video game reinforcement learning tasks. This preprocessing converts images to grayscale, downsamples spatial resolution to 84 × 84 pixels, and applies temporal downsampling to four frames while also computing the maximum pixel value between two consecutive frames to avoid flickering. The resulting preprocessed input is a (4 × 84 × 84) tensor.

The ResNet model takes raw frames as input, with temporal downsampling to 3.75 Hz to match the sampling frequency of the non-overlapping 16-frame window used in the base architecture.

### Models

All models in this study share the same base architecture, except for the pretrained ResNet baseline model. The base architecture consists of four convolutional layers followed by a fully connected layer (see **Figure 1**). The total parameter count of this shared base is 619,264. Each model includes an additional fully connected output layer, whose dimensionality depends on the number of outputs required by its training objective.

**Fig 1.**
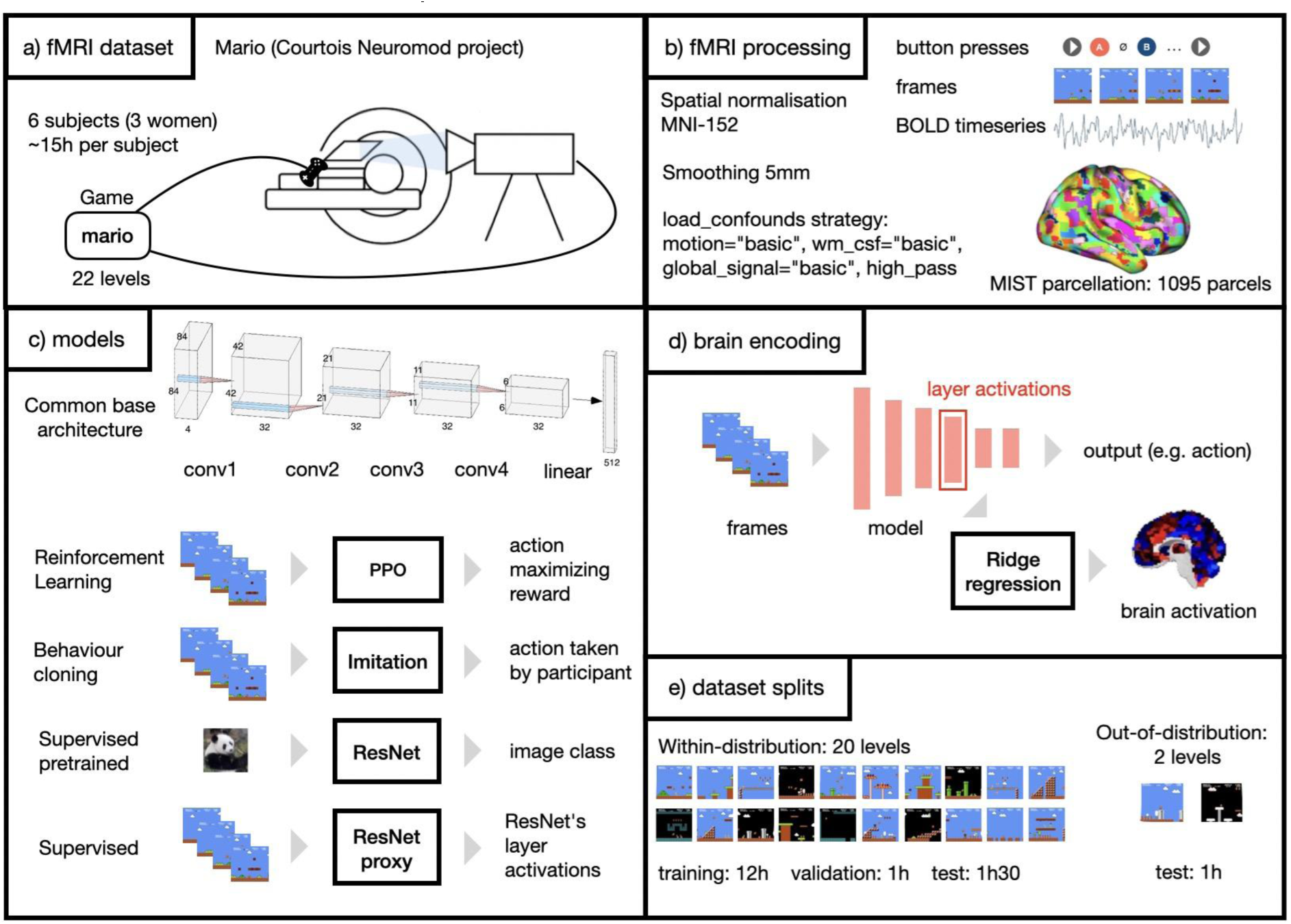
Overview of the study methods. fMRI data from subjects playing Super Mario Bros **(a)** is processed and reduced to time series with a brain parcellation **(b)**. Artificial models are trained with different objectives and tested for their ability to predict brain activity, using different objectives during training **(c)**: reinforcement learning, behavioural cloning and visual supervised learning (ResNet variants). Brain encoding performance **(d)** is evaluated both within-distribution (on the game levels used for training) and out-of-distribution (on game levels not seen during training), highlighting each model’s ability to generalize beyond their training data **(e)**.

#### PPO model

The PPO model follows a reinforcement learning procedure using the PPO algorithm (Schulman et al. 2017). The model’s objective is to maximize rewards while interacting with the game. The reward function incentivizes the agent to complete levels by rewarding rightward movement and score increases while penalizing time spent and losing a life. The output layer maps to a one-hot vector representing 12 possible actions. Training occurs across all 20 selected levels, with each rollout starting from a random position, ensuring exposure to all parts of the training levels. The total parameter count of the PPO model is 625,933.

#### Imitation models

Five Imitation models were trained, one per participant, using a supervised learning approach based on each player’s button presses. These models perform a *behavioral cloning* task, predicting which action the player took. The output layer maps to a binary vector representing which of six buttons were pressed. Each model is trained on video frames and button press data from its corresponding subject’s gameplay. Given a 16-frame sequence input, the model predicts the button presses in the next frame. The total parameter count of the Imitation models is 625,420.

#### ResNet model

The pretrained *ResNet152v2* model (He et al. 2015) is a convolutional neural network (CNN) with residual connections, trained on the *ImageNet1K* dataset (1.28 million images, 1,000 categories). This model is purely vision-based, trained to recognize object categories in natural images rather than video game frames. Unlike the other models, only a single layer of ResNet (the 145th) was used for brain encoding, as it was chosen to generate targets for the ResNet proxy model. This deeper layer was selected because it contains high-level semantic representations rather than low-level visual features. It is the only model in the comparison that is not sharing the same architecture and training data distribution as the other models. The total parameter count of the ResNet is 60,192,808.

#### ResNet proxy model

The *ResNet proxy model* was designed to replicate the representational space of the ResNet model while being trained on the same data distribution as the other models. It was trained in a supervised manner: for each 16-frame gameplay sequence, the final frame was passed through ResNet, and activations from its 145th layer were recorded. An incremental PCA was fitted on 20% of the training dataset to retain the top 1,000 principal components of these activations. The ResNet proxy model was then trained to predict these principal components. Using a proxy model instead of ResNet152v2 mitigates biases arising from differences in training datasets and architecture, ensuring a fairer comparison of brain encoding performance. The total parameter count of the ResNet proxy model is 1,132,264.

#### Untrained model

The *Untrained model* is a model with randomly initialized weights. It is not intended as an estimate of chance level, as it still processes game images. While applying random transformations to pixel values, it retains information from the images, producing arbitrary but structured representations. This model serves as a baseline to assess whether trained models produce representations more aligned with brain activity than random transformations of the same input.

### Brain encoding

For each model, subject, and layer, brain encoding was performed by extracting the activations from the layer of interest, convolving these activations with a model of the hemodynamic response function (HRF), and fitting a ridge regression to predict brain activity. The regression targets were the preprocessed BOLD time series, projected into parcels defined by the MIST atlas. Ridge regression was trained on each subject’s training set using a grid of alpha values (10^k^ with k ranging from -1 to 6). The optimal alpha was selected based on the lowest mean squared error (MSE) on the subject’s validation set. The brain encoding performance, reported in terms of the coefficient of determination (R²), was evaluated on either the test set or the out-of-distribution (held-out levels) set.

The HRF model used was the SPM canonical HRF as implemented in the nilearn Python library. Brain encoding was performed on all layers of each model, except the first and final layers. The first layer, due to its high dimensionality, was computationally expensive and yielded relatively poor encoding accuracy in preliminary analyses. The final layer was excluded because its dimensionality and function varied across models depending on their respective training objectives.

### Task performance metrics

#### Brain encoding

To assess how well each model predicts brain activity, we used the coefficient of determination (R²) between the original and predicted BOLD time series in each parcel of the MIST atlas. R² reflects the proportion of variance in the original signal explained by the predicted signal. A perfect prediction yields an R² score of 1, while a score of 0 indicates that the model does no better than predicting the mean signal. Negative R² values can occur when the prediction is worse than the mean, indicating uncorrelated or misleading variance in the prediction.

#### Game performance

To evaluate gameplay ability, each model was used to play each level 20 times for a maximum of 5000 frames (approximately 1 minute and 23 seconds). If the agent reached a game over before the time limit, it restarted with a new set of three lives. Since all levels are linear with the goal of progressing to the right, the maximum distance reached from the beginning of the level serves as a reliable proxy for task success. Game performance is reported as the average of the maximum distances reached across the 20 runs per level, averaged again across all levels.

#### Behavioral imitation

To measure how closely the models’ actions align with human behavior, we use a behavioral imitation score. This is computed by feeding the models sequences of frames from a subject’s gameplay recording, and comparing the predicted button presses to the actual button presses from the human player. The score is calculated as the average F1 score per button, averaged over buttons. The “up” and “down” buttons are excluded from this metric, as they are rarely used and not required to complete most levels.

### Regions of interest

To characterize the spatial distribution of brain encoding R² scores, we report not only parcel-wise maps but also scores averaged over functionally relevant regions of interest (ROIs).

Seven ROIs were derived from the Yeo7 network atlas (Thomas Yeo et al. 2011), corresponding to the following large-scale functional networks: visual, dorsal attention, sensorimotor, ventral attention, default mode, fronto-parietal, and limbic (see Supplementary Material Figure I).

The remaining five ROIs were defined using two vision localizer tasks completed in the scanner by three of the five participants (sub-01, sub-02, and sub-03). For sub-05 and sub-06, the regions derived from the data of sub-01 were used. The V1, V2, and V3 ROIs were identified using a retinotopy task adapted from Kay et al. (2013) and implemented in PsychoPy. Voxel-wise population receptive fields were estimated using the *analyzePRF* MATLAB toolbox (Kay et al. 2013), and subject-specific ROIs were refined using group atlas priors with *Neuropythy* (https://github.com/noahbenson/neuropythy) (Benson and Winawer 2018).

The fusiform face area (FFA) and parahippocampal place area (PPA) were identified using a PsychoPy implementation (https://github.com/NBCLab/pyfLoc) of the fLoc task developed by the Stanford VPN Lab (Stigliani et al. 2015), with stimuli sourced from the fLoc functional localizer package (https://github.com/VPNL/fLoc). ROI boundaries were defined from subject-specific contrasts, identifying voxels with preferential responses to specific stimulus categories. These contrasts were constrained within group-derived parcels of category-selective regions based on (Julian et al. 2012). The V1, V2, V3, FFA, and PPA ROIs for sub-01 are shown in **Figure 3**.

**Figure 2:**
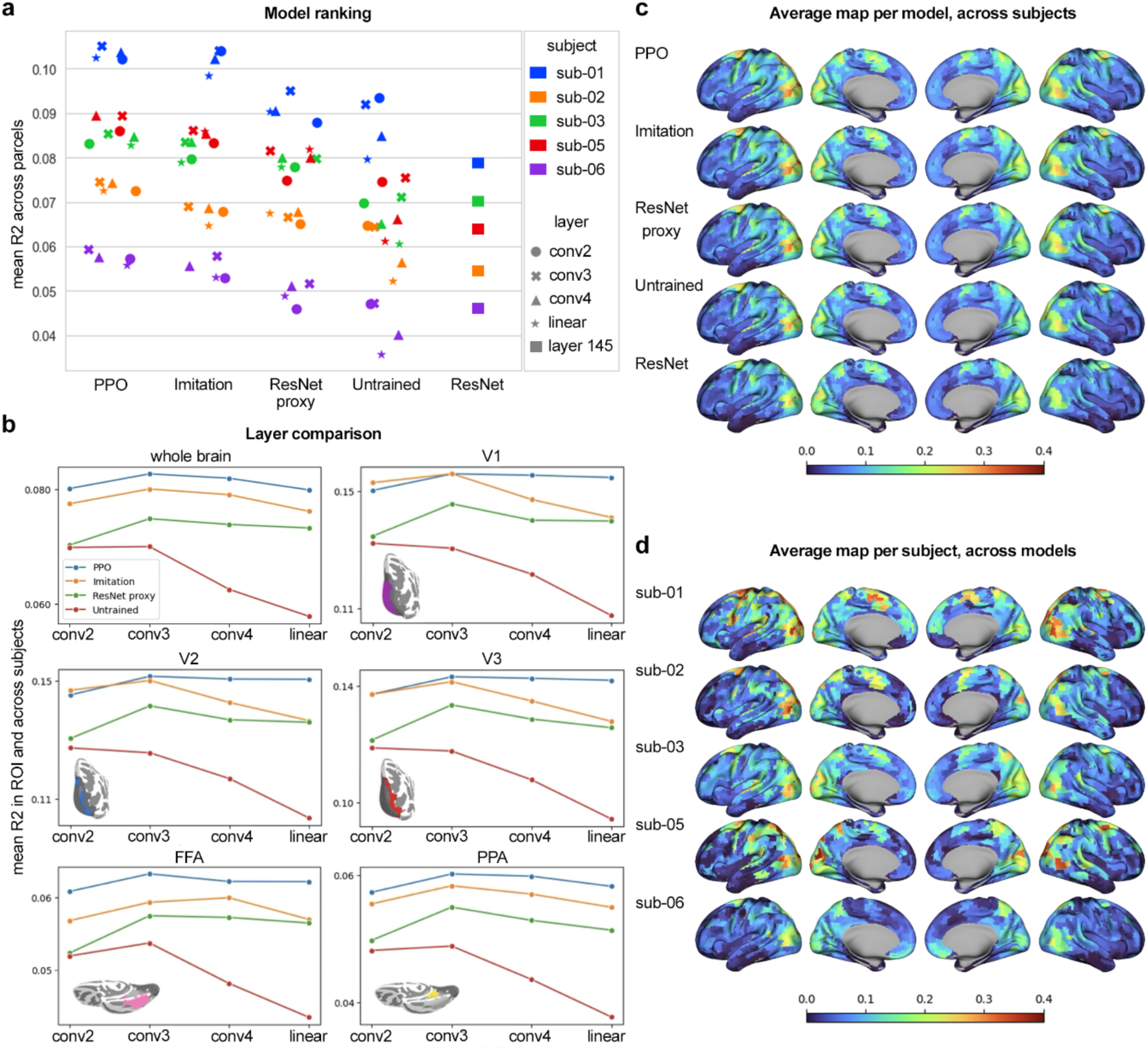
Comparison of the brain encoding R² scores of the different models. **a** Brain encoding R2 scores, averaged over brain parcels. Each dot corresponds to the value for a layer and a subject (the ResNet being evaluated on just one layer, it presents less sample points). **b** Mean brain encoding R² score per layer for each model. Each plot represents the scores averaged of the whole brain (top left) or different ROIs of the visual cortex. **c** Brain encoding maps per model. The maps represent the brain encoding R² score per parcel, averaged over subjects, for each model’s conv3 layer (except for the ResNet). For each model we see a similar spatial distribution of the R² score. The regions that get the highest R² scores are located in the visual cortex, the dorsal attentional path and in the motor cortex. **d** Brain encoding maps per subject. The maps represent the brain encoding R² score averaged over models, for the conv3 layer (except for the ResNet). The maps of the different subjects show more noticeable differences in R² amplitude than the maps per model, but the overall spatial distributions remain very similar. The regions that get the highest R² scores are located in the visual cortex, the dorsal attentional path and in the motor cortex.

**Figure 3:**
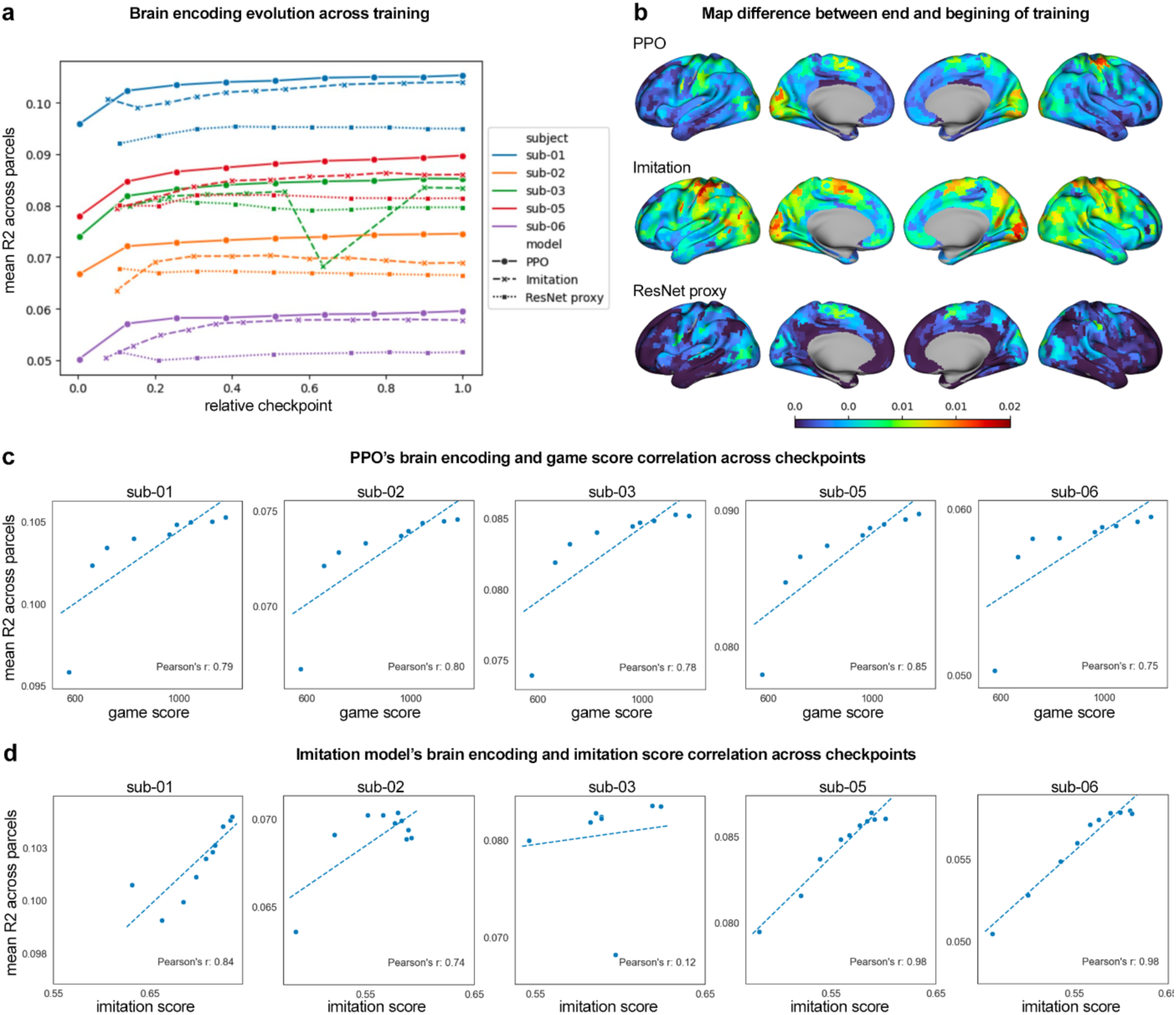
Training effects on brain encoding and task performance. **a**: Brain encoding R² for each model and subject across training checkpoints (averaged over brain parcels, evaluated on the test set). **b**: Difference maps showing changes in R² between early training (10–15%) and the final checkpoint, averaged across subjects. **c**: Correlation between brain encoding and game score for the PPO model across checkpoints. Dashed lines show linear fits. **d**: Correlation between brain encoding and imitation score for the Imitation models across checkpoints. Dashed lines show linear fits.

For each ROI, parcel-wise R² maps were projected back to MNI voxel space, and the ROI masks were applied to compute average R² values per region.

## Results

### Aim 1: ranking models through brain encoding performance

Our first aim was to rank models in terms of brain encoding performance, after completing training on within-distribution data. A separate brain encoding model was trained for each layer, across four layers (conv2, conv3, conv4, linear), using 1095 brain parcels as targets.

The PPO model achieved the highest overall brain encoding R², on average across subjects and model layer, see **Figure 2a**. The second-best model was the Imitation model, followed by the ResNet proxy model, and then the Untrained model. This ranking was statistically significant across nearly all layers and subjects (p<0.05, paired WIlcoxon two-sided test, Bonferroni-corrected; see **Table 1**). Yet, the differences in brain encoding accuracy across different learning objectives only had a limited magnitude, less than 1 percent of R².

**Table 1:**
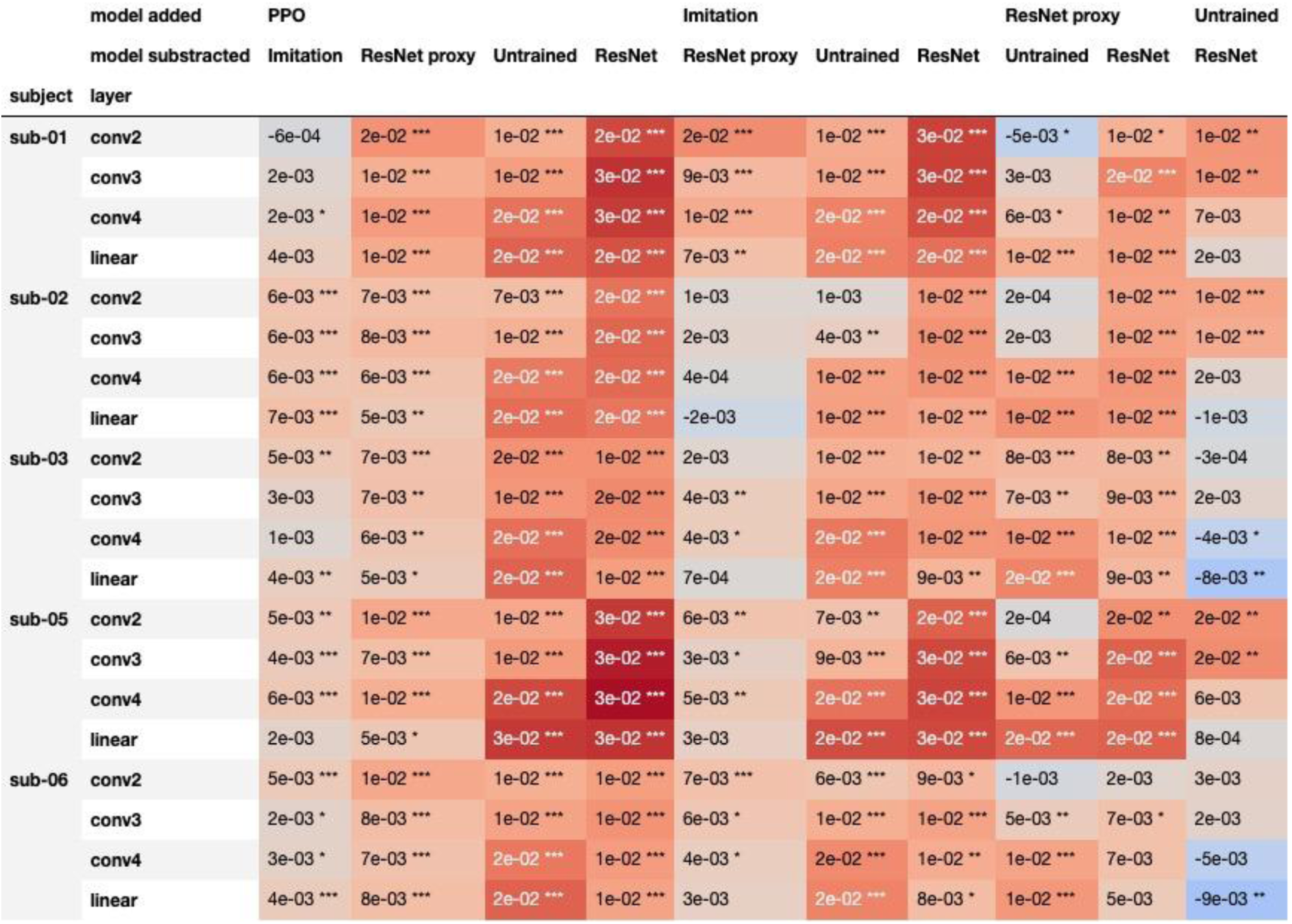
Difference of brain encoding R² between models for each subject and layer. The R² scores are averaged across brain parcels and across sequences of 100 TRs. The asterisks denote significance of a two sided Wilcoxon signed-rank test. One, two or three asterisks correspond respectively to a p-value < 0.05, a p-value FDR corrected (Benjamini/Yekutieli) < 0.05 and a p-value Bonferroni corrected < 0.05.

Interestingly, the brain encoding performance of the ResNet model was not significantly different from that of the Untrained model. The R² scores of the Untrained model were well above chance: a model trained on permuted input data has an R² very close to 0 (<0.001, results not shown). This indicated that even random projections of pixel-level data may be sufficient to encode brain activity in this video game task.

Overall, brain encoding allowed for a statistically significant ranking of model training objectives, though performance differences among the top models remained small.

### Aim 2: probing parallel hierarchies of representations between models and the brain

#### The conv3 layer has the highest brain encoding score, for trained models

We next compared the brain encoding accuracy between layers, for all the models sharing the same architecture. For each type of trained model (PPO, imitation and Resnet proxy), the 3rd convolutional layer of the models was the one producing the highest brain encoding accuracy, on average over subjects and brain parcels. The difference of R² across layers remained small in amplitude - less than .5%. This pattern remains consistent when averaging the R² scores on specific brain regions, notably in the visual cortex (see **Figure 2b**). By contrast, for the Untrained model, the accuracy of the conv3 layer was almost the same as conv2, and accuracy dropped sharply for subsequent levels after that.

Overall, the 3rd convolutional layer emerged as an optimal target for brain encoding and, for the following results, unless mentioned otherwise, the presented R² scores will correspond to this layer.

#### All the models (and layers) produced brain encoding maps with similar topography

Next, we examined whether different models encoded distinct brain regions. Within subjects, brain encoding maps were highly consistent across models, with an average Pearson’s spatial correlation coefficient of 0.97 (range: 0.92–0.99) across model pairs. This was evident in **Figure 2c**, which showed strikingly similar brain encoding maps for each model, averaged across subjects.

Across subjects, brain encoding maps for a given model were also similar but showed greater variability, with an average Pearson’s correlation coefficient of 0.62 (range: 0.50–0.67). This substantial individual variance is illustrated in **Figure 2d**, presenting each subject’s brain encoding maps, averaged across models.

The maps were also very consistent across layers, see Supplementary Material II. The average Pearson’s correlation coefficient between two maps of the same subject and model but different layers was 0.995 (with values ranging from 0.982 to 0.999).

The regions where the models could explain the highest proportion of the brain activity signal were the visual and the dorsal attentional networks, as defined by the Yeo-Krienen 7 resting-state network parcellation. Moreover, more variance was explained in the primary visual cortex (V1, V2 and V3) than in more specialized areas of the secondary visual cortex (FFA and PPA), see Supplementary Material III.

Overall, we found that the brain encoding maps were extremely consistent across models, capturing mostly visual and dorsal attentional cortices with substantial inter-subject variance.

### Aim 3: Relationship between task performance and brain encoding accuracy

We next examined whether model performance on the task (game score or imitation) was associated with its ability to encode individual brain activity.

#### Robust association between task performance and brain encoding across training stages

We first tested this relationship across different stages of training, as models gradually improved at the task. Model checkpoints saved at different stages revealed a steady increase in brain encoding accuracy (**Figure 3a**), with the exception of the ResNet proxy model, which showed little improvement. Gains were concentrated in the same regions that already showed strong encoding (**Figure 3b**).

We quantified PPO performance as the average distance travelled per level (**Figure 3c**). Although distinct from the RL reward signal, this measure similarly reflects the model’s ability to progress through the game. For all subjects, PPO brain encoding accuracy correlated strongly with this performance across training checkpoints. For the Imitation models, brain encoding accuracy tracked closely with their behavioural imitation score, which measures the similarity between model and subject actions on recorded gameplay (**Figure 3d**). Strong correlations (Pearson’s r > 0.7) were observed for all subjects except sub-03.

Together, these analyses show that task performance and brain encoding were linked during training, as brain encoding steadily improved as training progressed for both the PPO and imitation models.

#### Level patterns strongly predict brain encoding for fully trained models but not task performance

We examined next how task performance was associated with brain encoding accuracy, this time for fully trained models across different game levels. We first observed that task performance itself was only weakly associated with brain encoding accuracy across game levels. For the PPO models, correlations were weak or even negative (**Figure 4a**). For the Imitation models, correlations were also weak (0.15–0.27) in most subjects, with the exception of sub-02 (**Figure 4b**).

**Figure 4:**
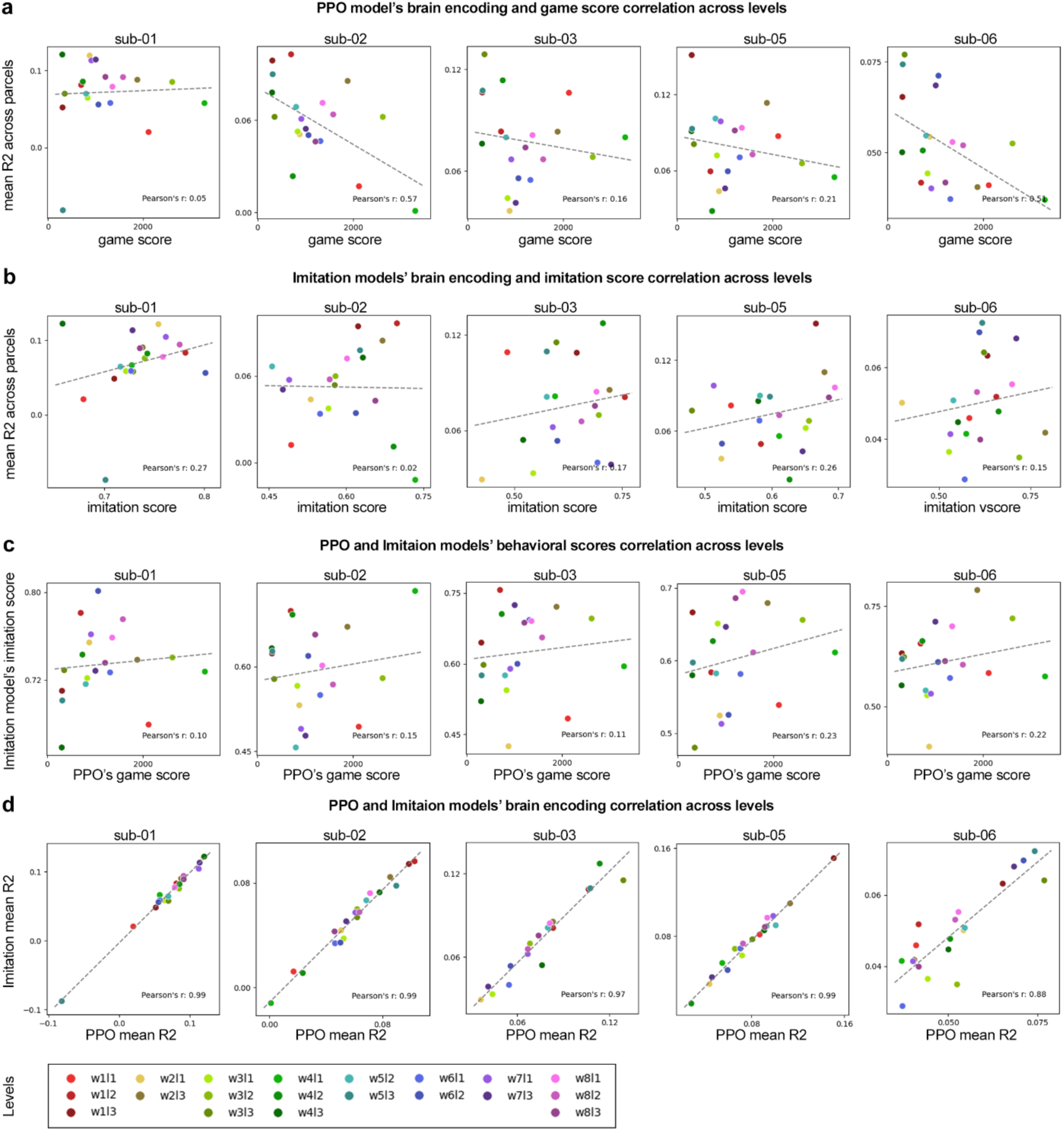
Cross-level correlations of brain encoding and task performance. **a**: PPO brain encoding versus game score across levels, shown separately for each subject. **b**: Imitation model brain encoding versus imitation score across levels, by subject. **c**: PPO game score versus imitation model score across levels, by subject. **d**: PPO and imitation model brain encoding across levels, showing near-perfect correlations.

To further investigate this lack of association, we compared the PPO and Imitation models across levels, separately in terms of performance or brain encoding (**Figure 4**), and a striking dichotomy emerged. PPO and Imitation models showed little correspondence in task performance (**Figure 4c**), suggesting that each faced different challenges. Yet their brain encoding scores were almost perfectly aligned (r = 0.88–0.99; **Figure 4d**). This pattern suggests that brain encoding quality was shaped more by the level itself than by model performance on that level.

### Aim 4: Out-of-distribution generalization of task performance and brain encoding

Our third aim was to assess how brain encoding and task performance generalize to out-of-distribution data. To do this, we measured game/imitation and brain encoding performance on two game levels excluded from training.

#### Training PPO models does not translate to out-of-distribution task performance, while the generalization of Imitation models was variable with clear benefits from training

In terms of game performance, for PPO, neither of the out-of-distribution levels benefited from training: the performance curves were flat across checkpoints, and fell outside of the distribution observed in the within-distribution level (**Figure 5a**, first graph). In terms of Imitation models, performance improved with training and fell within the within-distribution performance on one level (see orange curves **Figure 5a**), while the performance was lower than within-distribution on the other level and the effect of training was less pronounced (only apparent is sub-02, sub-05 and sub-06, see blue curves **Figure 5a**). Our results thus demonstrated that the behaviour of the PPO model is brittle, while the generalization of the Imitation models varied, although some benefits of training were generally observed.

**Figure 5:**
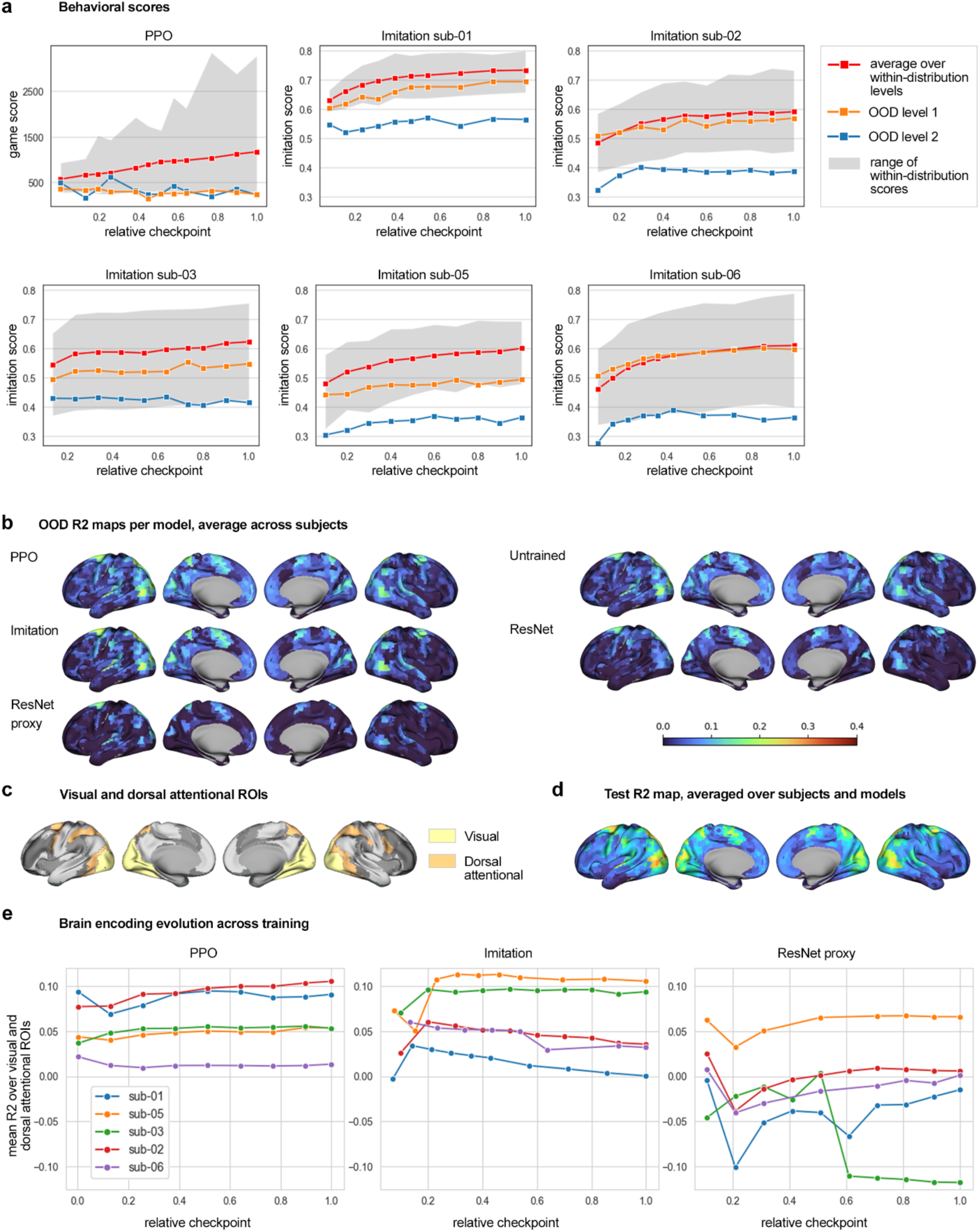
Out-of-distribution evaluation. **a** Behavioral scores comparison between the average on within-distribution levels (red) and the scores and each of the OOD levels (orange and blue). For the PPO, the behavioral score is the game performance score. For the Imitation models, the score is the imitation score. **b** Brain encoding maps of the OOD levels, for each model (conv3 layer for all models but ResNet) and averaged over subjects. **c** Visual and dorsal attentional ROIs from the Yeo7 atlas. **d** Brain encoding map for the within-distribution levels, averaged over subjects and models. **e** Evolution of the brain encoding accuracy across training in the visual and dorsal attentional ROIs.

#### Brain encoding performance was severely degraded out-of-distribution for all models

Across all models, R² scores dropped substantially on out-of-distribution levels (**Figure 5b**). This decline was particularly severe for the ResNet proxy model, which frequently produced negative R² values, indicating a complete failure to generalize. Differences between models became more pronounced in this setting: PPO exhibited the best generalization on average, followed by the Untrained model, then the Imitation models. Regarding spatial distribution of R² scores, the visual and dorsal attention networks showed the highest generalization on average, which were already networks with high within-distribution encoding performance, see **Figure 5b**. The lack of generalization in game/imitation performance of the PPO and imitation alone does not explain the out-of-distribution collapse in brain encoding entirely, as similar failures were observed for both the Untrained model and the pretrained ResNet. This shows that the ridge regression layer itself lacked generalization ability, rather than the failure being solely due to the model’s internal representations.

#### Effect of training on brain encoding was apparent for PPO and Imitation models

PPO models evaluated at different training checkpoints showed small improvement in brain encoding performance, after an initial dip in performance (**Figure 5c**, left graph). The Imitation models benefitted more uniformly from the earliest phase of training (**Figure 5c**, middle graph). Finally, the behaviour of the ResNet proxy showed R2 in the negative range for most of the training for all subjects but sub-05, thus indicating an overall failure to learn generalizable features (**Figure 5c**, right graph).

Overall, we found that our within-distribution findings regarding game/imitation and brain encoding performance did not translate to out-of-distribution levels. This demonstrates that the models we tested are limited in scope and may not provide a valid inference of brain-like processing, as human behavior remains robust and generalizable across levels. In this sense, all tested models appeared misaligned with brain data when evaluated on out-of-distribution tasks.

## Discussion

### Aim 1: A consistent ranking was observed across models, yet only with small absolute differences in brain encoding performance

As expected we observed that the action models consistently outperformed the vision and untrained baselines. Although some prior works had hypothesized that RL (Cross et al. 2021) or imitation (Kemtur et al. 2023) would lead to a better brain encoding, these works had not performed well controlled experiments which would have allowed us to make a specific prediction for our experiments regarding the RL vs imitation. The RL (PPO) objective appeared to have superior performance in brain encoding, consistently across subjects.

Surprisingly, the absolute differences in the quality of brain encoding between models were rather small. In particular, even the Untrained model was able to encode brain activity with a performance that approached that of our best models. This is not something that is typically observed in brain encoding of complex stimuli, such as natural images or videos. For instance, the 2025 Algonauts competition on video watching implemented a competent baseline of vision-language-audio processing, yet the best models explained up to twice as much variance as the baseline, and this difference was even more dramatic in out-of-distribution experiments (Gifford et al. 2025). However, in our set up, the diversity of visual stimuli was very limited, as it was constrained by the sprites of the video game. The same sprites were used in the train and test datasets, even though the gameplay may have been different, and this limitation even applied to out-of-distribution experiments to a large degree. This means that high-level abstract events, such as a jump, can be detected by brittle features encoding a specific combination of pixel values, which an Untrained network can represent well. We thus speculate that videogames are particularly prone to brittle brain encoding, at least when the exact same game mechanics are used during training and testing. This overfitting caveat had already been identified in the field of RL where agents are also commonly trained and tested in the same environment (Henderson et al. 2019).

### Aim 2: No hierarchy of brain encoding maps emerged from either the training objective or the target layer

We had hypothesized a hierarchy in the quality of brain encoding that would be related to the nature of the training objective: ResNet and ResNet proxy models would produce better encoding in the visual cortex, while the PPO and Imitation models would encode better in the sensorimotor cortex, as they were trained to perform actions in the game. This hypothesis was supported by recent results comparing monomodal vs multimodal vision-language-auditory representations during video watching, where models trained with a monomodal objective encoded better the specific brain network matching their training modality, yet multimodal representation encoded better association cortices (d’Ascoli et al. 2025). We also expected the Untrained model, with no task objective, to markedly underperform compared to trained models in task-relevant regions. Our results did not support our hypothesis. This may seem particularly at odds with the results reported by Cross and colleagues (Cross et al. 2021) who had reported a hierarchy of representations comparing a vision and action model in classic Atari games. However Cross and colleagues did not compare the same layer activations of their RL agents and vision baseline, using a concatenation of all layers for the RL models and only the last layer of the encoder for the vision model. This difference in activation sampling may explain this discrepancy in results with our study. In our case, we found that the intermediate layer was best for encoding brain activity irrespective of the training objective, and that when using this optimal layer there was no clear hierarchy of representations between training objectives.

We had also hypothesized a hierarchical correspondence between model depth and cortical areas. Such correspondences have been reported for some visual models (Nonaka et al. 2021), but recent work has shown that hierarchical structure is not necessary for convolutional neural networks (CNNs) to capture activity across visual regions (St-Yves et al. 2023). Contrary to our hypothesis, we observed that the third convolutional layer consistently produced the highest brain encoding accuracy. This pattern held across brain regions, including the visual cortex, and did not support a hierarchical mapping between network layer depth and brain regions. This is inconsistent with results reported by (Kemtur et al. 2023), which may reflect the absence of recurrence in our tested models. Recurrent layers were prominently featured in the architectures of Kemtur and colleagues and drove layer differentiation.

### Aim 3: Performance on the training task correlates with brain encoding accuracy, but systematic gaps in brain encoding performance were observed across levels

For each model individually, we found a correlation between task performance and brain encoding accuracy across training checkpoints, mirroring results from language on both fMRI and EEG data (Caucheteux and King 2022) and vision on intracranial neural recordings (Yamins et al. 2014). Still, these improvements were modest in amplitude and did not reflect qualitative differences in the topography of brain encoding maps across training objectives, contrary to our initial hypotheses. By contrast we found a massive gap in brain encoding performance across levels, that correlated highly between models trained with PPO and imitation. Our findings align with recent work by Conwell and colleagues (Conwell et al. 2023), who found that apparent differences in brain encoding across training tasks often reflect underlying differences in training datasets, particularly their diversity, more than the objective function itself. Wang and colleagues similarly emphasize the role of data diversity in shaping brain-model alignment (Wang et al. 2023).

### Aim 4: Out-of-distribution evaluation reveals brittle generalization of brain encoding

All models showed a marked decrease in brain encoding accuracy when evaluated on out-of-distribution (OOD) data. Differences between models actually became more pronounced under OOD conditions. PPO, Imitation, and Untrained models outperformed the ResNet-based vision models, with the ResNet proxy—competitive under in-distribution testing—collapsing to near-chance on OOD data. PPO models generalized better than the Untrained baseline, while Imitation models, although weaker overall, outperformed it in task-relevant areas such as V1, V2, and the dorsal attention network. So while all models demonstrated brittle performance, the *extent* of their brittleness revealed critical differences across training objectives.

### Limitations and future work

It should be emphasized that a model’s ability to perform brain encoding does not, by itself, imply functional similarity to the brain. To strengthen the interpretability of brain encoding results, we believe it is essential to evaluate predictive power across a wide range of stimuli and to compare performance across diverse data distributions, a point we previously demonstrated by showing the memory content of the videogame emulator itself could encode brain activity in Mario (Harel, Paugam, et al. 2025), but in a very brittle fashion. In this work, we assessed out-of-distribution (OOD) performance using two left-out game levels and observed that these levels behaved quite differently. The small number and arbitrary selection of these levels make it impossible to determine which specific factors contributed to the observed OOD performance. Future studies using videogame environments could systematically probe a larger and more controlled variety of OOD levels (or sublevels) to identify which task, visual, or structural features most strongly impact generalization performance, and we recently proposed a framework to implement such experiments by segmenting levels into short scenes featuring well identified game patterns (Harel, Bellec, et al. 2025).

We finally believe that generalist agents will be required in order to achieve robust brain encoding that generalize across many environments. A current trend in AI research is the development of foundation models for vision-action instead of training models from scratch on a particular task, although these models have not yet become as successful and widely available as large language models. Alternatively, task-specific agents trained in the latent space of a vision foundation model (Zhou et al. 2024) may also show improved robustness compared to training agents from scratch.

## Conclusion

We evaluated how the training objective of a videogame model impacts its ability to encode brain activity. To do so, we rigorously controlled for potential differences related to architecture design and training data. Evaluations on 20 levels of the Super Mario Bros videogame revealed significant differences between models, with reinforcement learning performing best, though only with modest gains over imitation learning or even an untrained model. Contrary to our expectations based on prior literature, no model layers or specific training objectives preferentially aligned with specific brain areas. Although brain encoding accuracy steadily improved during training, large gaps persisted across game levels, even for fully trained models. Out-of-distribution experiments also showed brittle brain encoding performance, with poor generalization to new game levels.

Overall, our results show that small convolutional models are limited in their capacity to robustly encode brain activity, regardless of the training objective. We also found that diverse test data are essential to revealing these gaps in generalization, both within- and out-of-distribution. High-quality brain encoding for videogame tasks will likely require larger models with robust task performance, such as vision-action foundation models.

## BIBLIOGRAPHY

Ahmadi, Sana, Francois Paugam, Tristan Glatard, and Pierre Lune Bellec. 2024. “Training Compute-Optimal Vision Transformers for Brain Encoding.” arXiv:2410.19810. Preprint, arXiv, October 17. 10.48550/arXiv.2410.19810.

Allen, Emily J., Ghislain St-Yves, Yihan Wu, et al. 2022. “A Massive 7T fMRI Dataset to Bridge Cognitive Neuroscience and Artificial Intelligence.” Nature Neuroscience 25 (1): 116–26. 10.1038/s41593-021-00962-x.

Ascoli, Stéphane d’, Jérémy Rapin, Yohann Benchetrit, Hubert Banville, and Jean-Rémi King. 2025. “TRIBE: TRImodal Brain Encoder for Whole-Brain fMRI Response Prediction.” arXiv:2507.22229. Preprint, arXiv, July 29. 10.48550/arXiv.2507.22229.

Bellec, Pierre, and Julie Boyle. 2019. “Bridging the Gap between Perception and Action: The Case for Neuroimaging, AI and Video Games.” Preprint, OSF, July 5. 10.31234/osf.io/3epws.

Benson, Noah C, and Jonathan Winawer. 2018. “Bayesian Analysis of Retinotopic Maps.” eLife 7 (December): e40224. 10.7554/eLife.40224.

Boyle, Julie A., Basile Pinsard, Amal Boukhdhir, et al. 2020. “The courtois project on neuronal modelling - first data release.” Paper presented at 26th OHBM annual meeting. https://publications.polymtl.ca/50613/.

Brockman, Greg, Vicki Cheung, Ludwig Pettersson, et al. 2016. “OpenAI Gym.” arXiv:1606.01540. Preprint, arXiv, June 5. 10.48550/arXiv.1606.01540.

Burelli, Paolo, and Laurits Dixen. 2024. “Playing With Neuroscience: Past, Present and Future of Neuroimaging and Games.” arXiv:2403.15413. Preprint, arXiv, March 6. 10.48550/arXiv.2403.15413.

Caucheteux, Charlotte, and Jean-Rémi King. 2022. “Brains and Algorithms Partially Converge in Natural Language Processing.” Communications Biology 5 (1): 1. 10.1038/s42003-022-03036-1.

Cichy, Radoslaw Martin, Aditya Khosla, Dimitrios Pantazis, Antonio Torralba, and Aude Oliva. 2016. “Comparison of Deep Neural Networks to Spatio-Temporal Cortical Dynamics of Human Visual Object Recognition Reveals Hierarchical Correspondence.” Scientific Reports 6 (June): 27755. 10.1038/srep27755.

Conwell, Colin, Jacob S. Prince, Kendrick N. Kay, George A. Alvarez, and Talia Konkle. 2023. “What Can 1.8 Billion Regressions Tell Us about the Pressures Shaping High-Level Visual Representation in Brains and Machines?” Preprint, bioRxiv, July 1. 10.1101/2022.03.28.485868.

Cross, Logan, Jeff Cockburn, Yisong Yue, and John P. O’Doherty. 2021. “Using Deep Reinforcement Learning to Reveal How the Brain Encodes Abstract State-Space Representations in High-Dimensional Environments.” Neuron 109 (4): 724–738.e7. 10.1016/j.neuron.2020.11.021.

Deng, Jia, Wei Dong, Richard Socher, Li-Jia Li, Kai Li, and Li Fei-Fei. 2009. “ImageNet: A Large-Scale Hierarchical Image Database.” 2009 IEEE Conference on Computer Vision and Pattern Recognition, June, 248–55. 10.1109/CVPR.2009.5206848.

Eickenberg, Michael, Alexandre Gramfort, Gaël Varoquaux, and Bertrand Thirion. 2017. “Seeing It All: Convolutional Network Layers Map the Function of the Human Visual System.” NeuroImage 152 (May): 184–94. 10.1016/j.neuroimage.2016.10.001.

Freteault, Maelle, Maximilien Le Clei, Loic Tetrel, Pierre Bellec, and Nicolas Farrugia. 2023. Alignment of Auditory Artificial Networks with Massive Individual fMRI Brain Data Leads to Generalizable Improvements in Brain Encoding and Downstream Tasks. September 6. 10.1101/2023.09.06.556533.

Gifford, Alessandro T., Domenic Bersch, Marie St-Laurent, et al. 2025. “The Algonauts Project 2025 Challenge: How the Human Brain Makes Sense of Multimodal Movies.” arXiv:2501.00504. Preprint, arXiv, January 6. 10.48550/arXiv.2501.00504.

Güçlü, Umut, and Marcel A. J. van Gerven. 2015. “Deep Neural Networks Reveal a Gradient in the Complexity of Neural Representations across the Ventral Stream.” Articles. Journal of Neuroscience 35 (27): 10005–14. 10.1523/JNEUROSCI.5023-14.2015.

Harel, Yann, Lune P. Bellec, François Paugam, Hugo Delhaye, and Audrey Durand. 2025. “Human-AI Alignment of Learning Trajectories in Video Games: A Continual RL Benchmark Proposal.” Paper presented at Reinforcement Learning and Video Games Workshop @ RLC 2025. July 11. https://openreview.net/forum?id=YAVB439L9X.

Harel, Yann, François Paugam, Marie St-Laurent, and Pierre Bellec. 2025. “Brittle Brain Encoding: Poor Out-of-Distribution Generalization Shows the Human Brain Is Neither a Nintendo Entertainment System nor a Four-Layer Convolutional Neural Network.” Paper presented at Cognitive Computational Neuroscience, Amsterdam, Netherlands. August 12. https://2025.ccneuro.org/abstract_pdf/Harel_2025_Brittle_Brain_Encoding_Poor_Out-of-Distribution_Generalization.pdf.

Harel, Yann, Basile Pinsard, Julie Boyle, et al. 2023. “Gamer in the Scanner : Event-Related Analysis of fMRI Activity during Retro Videogame Play Guided by Automated Annotations of Game Content.” Preprint, OSF, October 27. 10.31234/osf.io/uakq9.

He, Kaiming, Xiangyu Zhang, Shaoqing Ren, and Jian Sun. 2015. “Deep Residual Learning for Image Recognition.” arXiv:1512.03385. Preprint, arXiv, December 10. 10.48550/arXiv.1512.03385.

Henderson, Peter, Riashat Islam, Philip Bachman, Joelle Pineau, Doina Precup, and David Meger. 2019. “Deep Reinforcement Learning That Matters.” arXiv:1709.06560. Preprint, arXiv, January 30. 10.48550/arXiv.1709.06560.

Jiang, Minqi, Michael Dennis, Jack Parker-Holder, Jakob Foerster, Edward Grefenstette, and Tim Rocktäschel. 2022. “Replay-Guided Adversarial Environment Design.” arXiv:2110.02439. Preprint, arXiv, January 17. 10.48550/arXiv.2110.02439.

Julian, J. B., Evelina Fedorenko, Jason Webster, and Nancy Kanwisher. 2012. “An Algorithmic Method for Functionally Defining Regions of Interest in the Ventral Visual Pathway.” NeuroImage 60 (4): 2357–64. 10.1016/j.neuroimage.2012.02.055.

Kay, Kendrick N., Jonathan Winawer, Aviv Mezer, and Brian A. Wandell. 2013. “Compressive Spatial Summation in Human Visual Cortex.” Journal of Neurophysiology 110 (2): 481. 10.1152/jn.00105.2013.

Kell, Alexander J. E., Daniel L. K. Yamins, Erica N. Shook, Sam V. Norman-Haignere, and Josh H. McDermott. 2018. “A Task-Optimized Neural Network Replicates Human Auditory Behavior, Predicts Brain Responses, and Reveals a Cortical Processing Hierarchy.” Neuron 98 (3): 630–644.e16. 10.1016/j.neuron.2018.03.044.

Kemtur, Anirudha, Francois Paugam, Basile Pinsard, et al. 2023. “Behavioral Imitation with Artificial Neural Networks Leads to Personalized Models of Brain Dynamics During Videogame Play.” Preprint, bioRxiv, November 29. 10.1101/2023.10.28.564546.

Mnih, Volodymyr, Koray Kavukcuoglu, David Silver, et al. 2015. “Human-Level Control through Deep Reinforcement Learning.” Nature 518 (7540): 7540. 10.1038/nature14236.

Nichol, Alex, Vicki Pfau, Christopher Hesse, Oleg Klimov, and John Schulman. 2018. “Gotta Learn Fast: A New Benchmark for Generalization in RL.” arXiv:1804.03720. Preprint, arXiv, April 23. 10.48550/arXiv.1804.03720.

Nonaka, Soma, Kei Majima, Shuntaro C. Aoki, and Yukiyasu Kamitani. 2021. “Brain Hierarchy Score: Which Deep Neural Networks Are Hierarchically Brain-Like?” iScience 24 (9): 103013. 10.1016/j.isci.2021.103013.

Paolo, Giuseppe, Jonas Gonzalez-Billandon, and Balázs Kégl. 2024. “A Call for Embodied AI.” arXiv:2402.03824. Preprint, arXiv, September 13. 10.48550/arXiv.2402.03824.

Paugam, François, Guillaume Lajoie, and Pierre Bellec. 2024. “What Is a Good Model for Brain Encoding in a Videogame Task ?” Paper presented at Computational Cognitive Neuroscience 2024, Boston, MA. August 9. https://2024.ccneuro.org/poster/?id=389.

Schrimpf, Martin, Idan Blank, Greta Tuckute, et al. 2020. “Artificial Neural Networks Accurately Predict Language Processing in the Brain.” Preprint, bioRxiv, June 27. 10.1101/2020.06.26.174482.

Schrimpf, Martin, Jonas Kubilius, Ha Hong, et al. 2018. “Brain-Score: Which Artificial Neural Network for Object Recognition Is Most Brain-Like?” Preprint, bioRxiv, September 5. 10.1101/407007.

Schulman, John, Filip Wolski, Prafulla Dhariwal, Alec Radford, and Oleg Klimov. 2017. “Proximal Policy Optimization Algorithms.” arXiv:1707.06347. Preprint, arXiv, August 28. 10.48550/arXiv.1707.06347.

Seeliger, K., L. Ambrogioni, Y. Güçlütürk, L. M. van den Bulk, U. Güçlü, and M. A. J. van Gerven. 2021. “End-to-End Neural System Identification with Neural Information Flow.” PLOS Computational Biology 17 (2): e1008558. 10.1371/journal.pcbi.1008558.

Stigliani, Anthony, Kevin S. Weiner, and Kalanit Grill-Spector. 2015. “Temporal Processing Capacity in High-Level Visual Cortex Is Domain Specific.” The Journal of Neuroscience: The Official Journal of the Society for Neuroscience 35 (36): 12412–24. 10.1523/JNEUROSCI.4822-14.2015.

St-Yves, Ghislain, Emily J. Allen, Yihan Wu, Kendrick Kay, and Thomas Naselaris. 2023. “Brain-Optimized Deep Neural Network Models of Human Visual Areas Learn Non-Hierarchical Representations.” Nature Communications 14 (1): 1. 10.1038/s41467-023-38674-4.

Thomas Yeo, B. T., Fenna M. Krienen, Jorge Sepulcre, et al. 2011. “The Organization of the Human Cerebral Cortex Estimated by Intrinsic Functional Connectivity.” Journal of Neurophysiology 106 (3): 1125–65. 10.1152/jn.00338.2011.

Toneva, Mariya, and Leila Wehbe. 2019. “Interpreting and Improving Natural-Language Processing (in Machines) with Natural Language-Processing (in the Brain).” arXiv:1905.11833. Preprint, arXiv, November 13. 10.48550/arXiv.1905.11833.

Wang, Aria Y., Kendrick Kay, Thomas Naselaris, Michael J. Tarr, and Leila Wehbe. 2023. “Natural Language Supervision with a Large and Diverse Dataset Builds Better Models of Human High-Level Visual Cortex.” Preprint, bioRxiv, July 11. 10.1101/2022.09.27.508760.

Yamins, Daniel L. K., and James J. DiCarlo. 2016. “Using Goal-Driven Deep Learning Models to Understand Sensory Cortex.” Nature Neuroscience 19 (3): 3. 10.1038/nn.4244.

Yamins, Daniel L. K., Ha Hong, Charles F. Cadieu, Ethan A. Solomon, Darren Seibert, and James J. DiCarlo. 2014. “Performance-Optimized Hierarchical Models Predict Neural Responses in Higher Visual Cortex.” Proceedings of the National Academy of Sciences 111 (23): 8619–24. 10.1073/pnas.1403112111.

Zador, Anthony, Sean Escola, Blake Richards, et al. 2023. “Catalyzing Next-Generation Artificial Intelligence through NeuroAI.” Nature Communications 14 (1): 1597. 10.1038/s41467-023-37180-x.

Zhou, Gaoyue, Hengkai Pan, Yann LeCun, and Lerrel Pinto. 2024. “DINO-WM: World Models on Pre-Trained Visual Features Enable Zero-Shot Planning.” arXiv:2411.04983. Preprint, arXiv, November 7. 10.48550/arXiv.2411.04983. ary Material

