## Supplementary Material for "Training neural networks from scratch in a videogame leads to brittle brain encoding"

Supplementary I : Yeo7 atlas

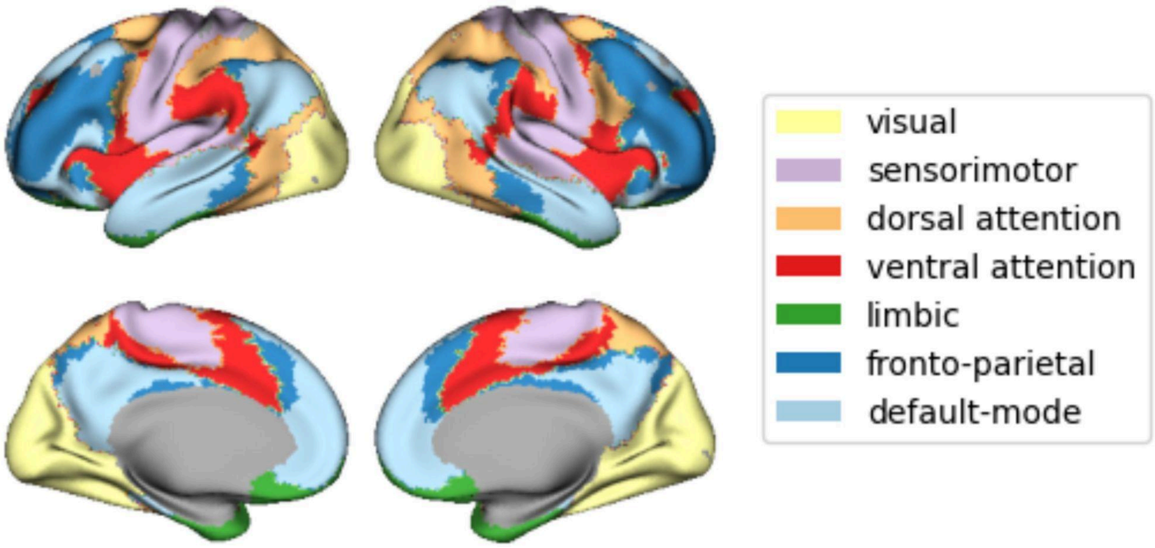


Visualization of the Yeo7 network atlas used to define regions of interest.

Supplementary II.i : PPO model brain maps

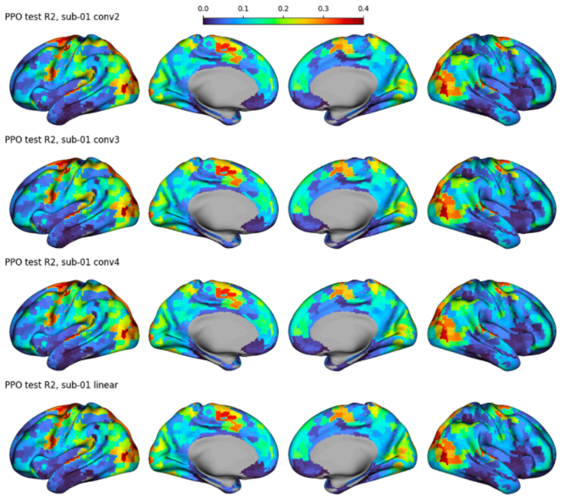


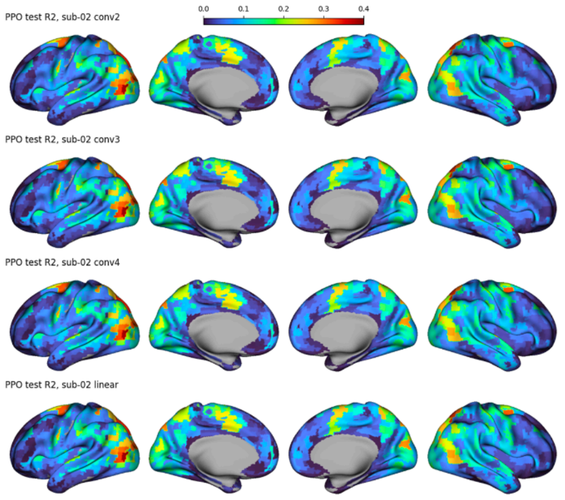


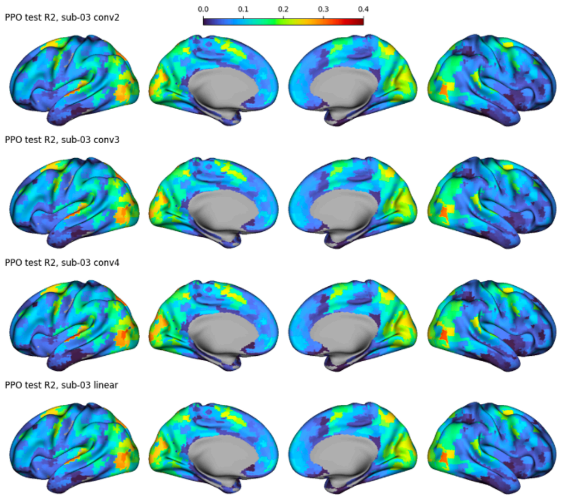


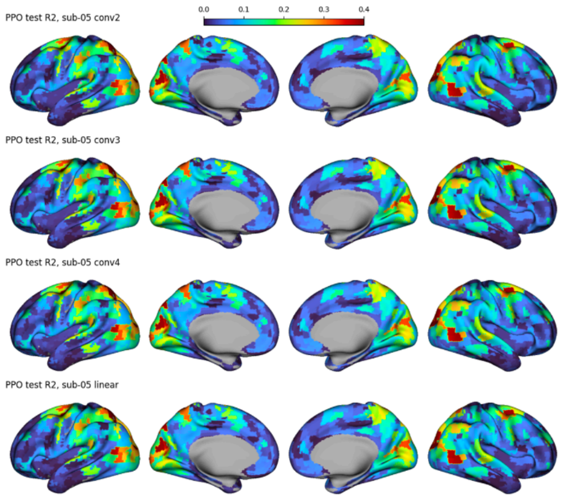


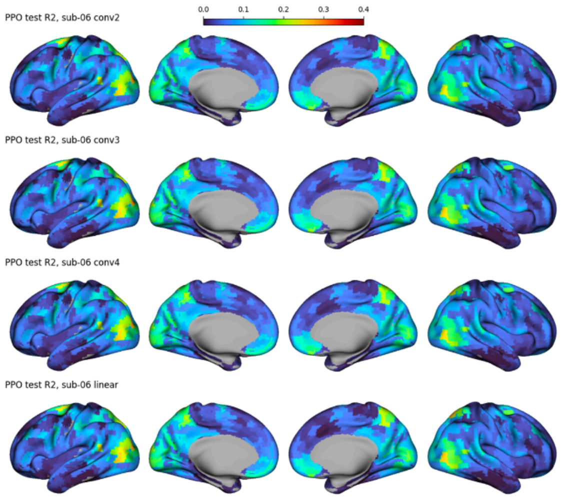

### Supplementary II.ii : Imitation model brain maps

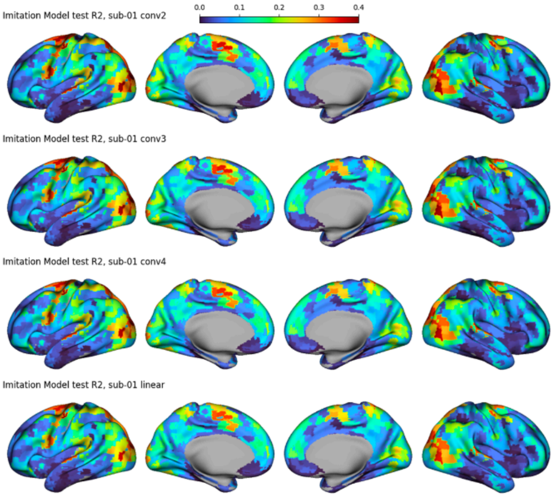


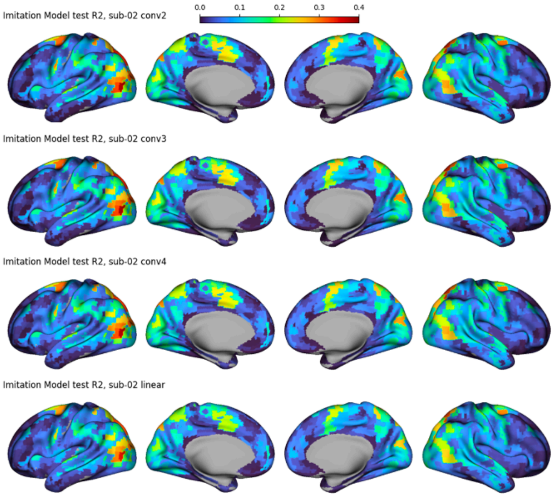


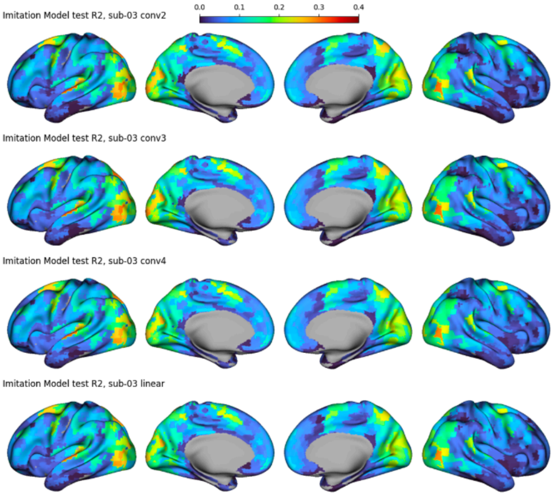


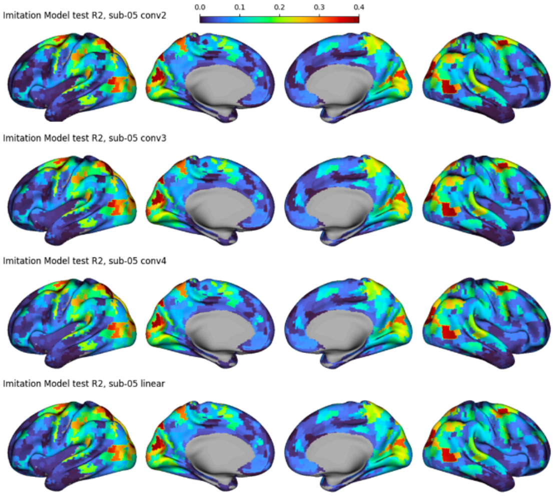


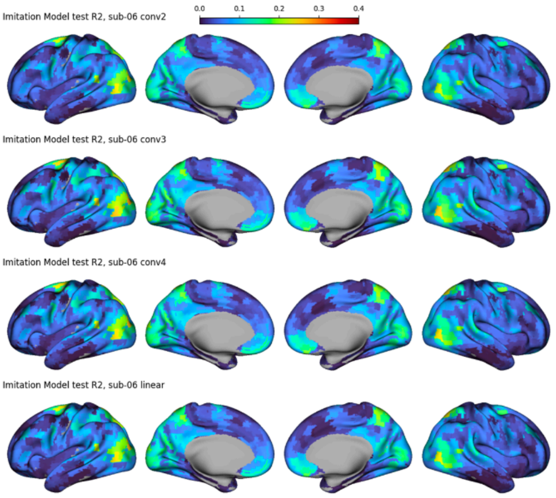

### Supplementary II.iii : ResNet proxy model brain maps

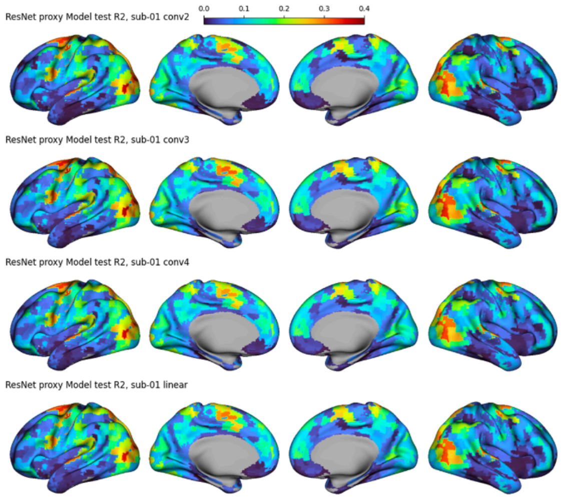


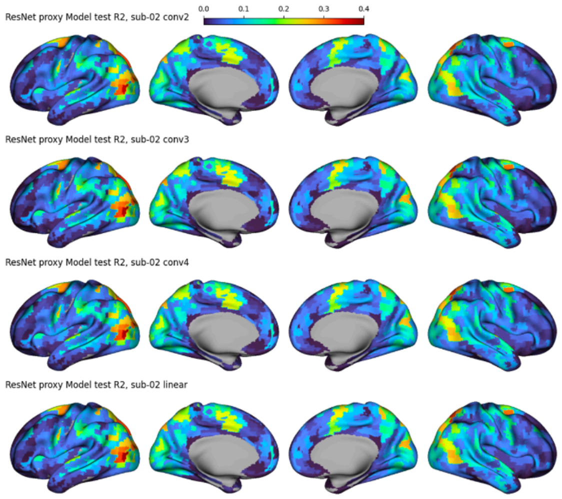


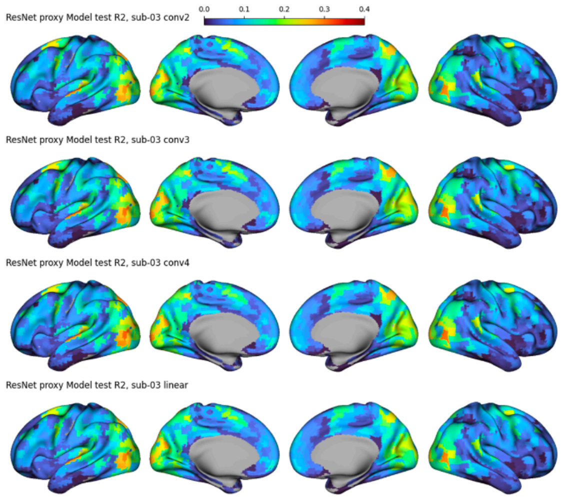


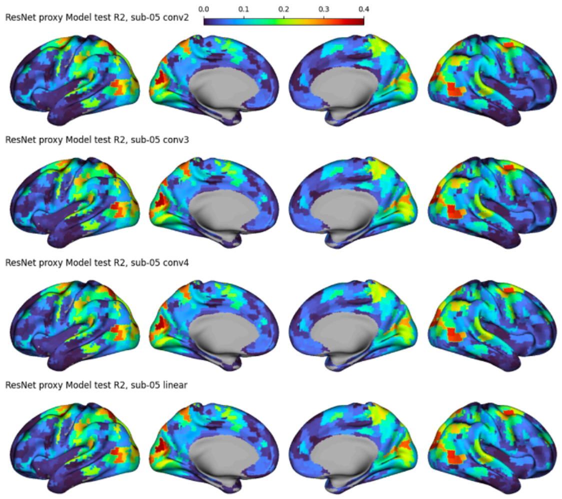


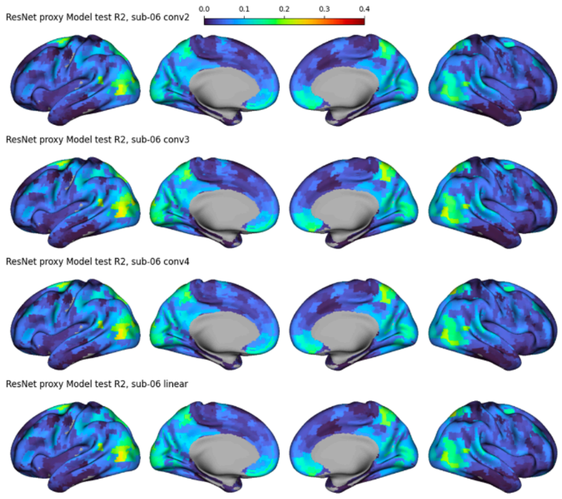

### Supplementary II.iv : Untrained model brain maps

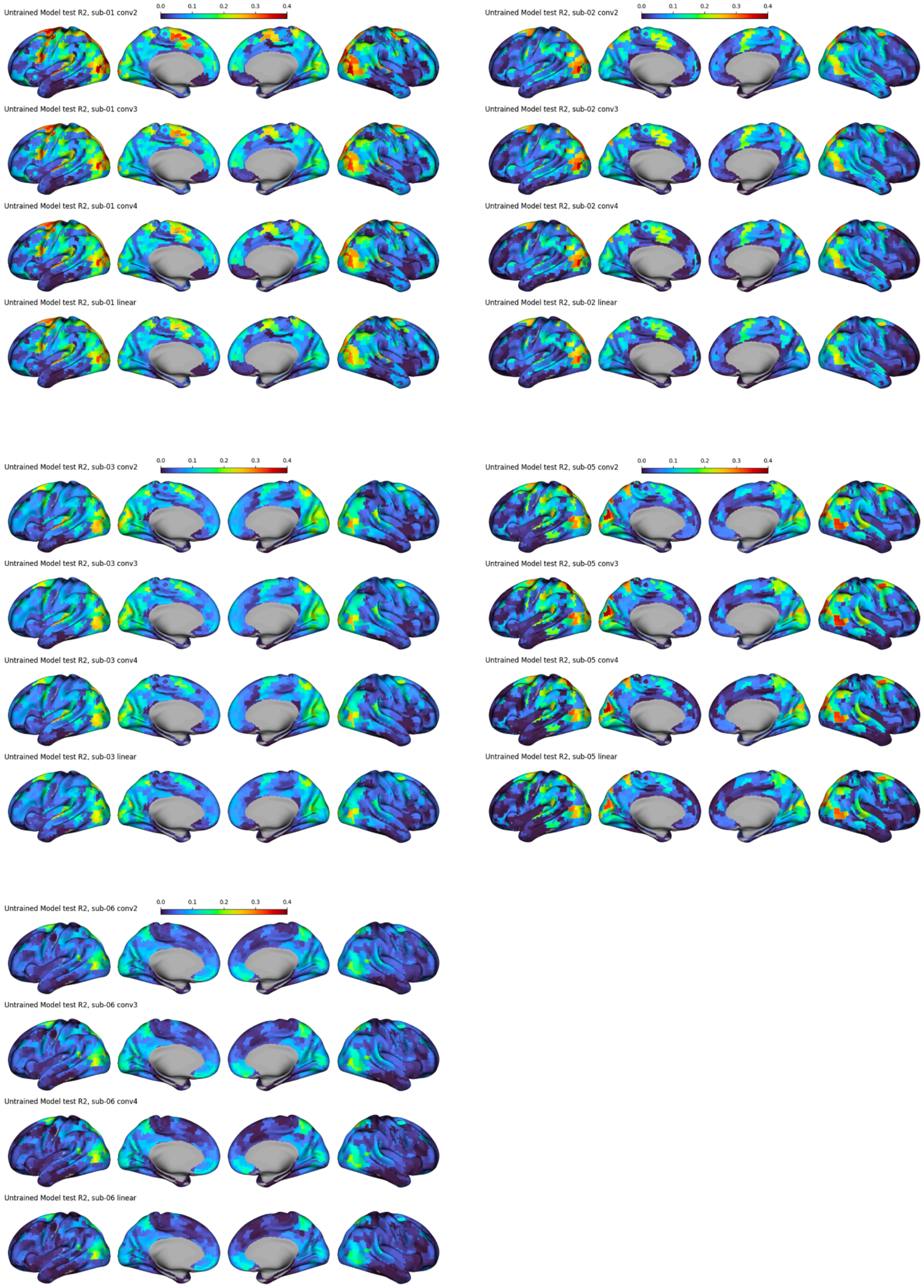

#### Supplementary III : Test brain encoding R2 per region of interest

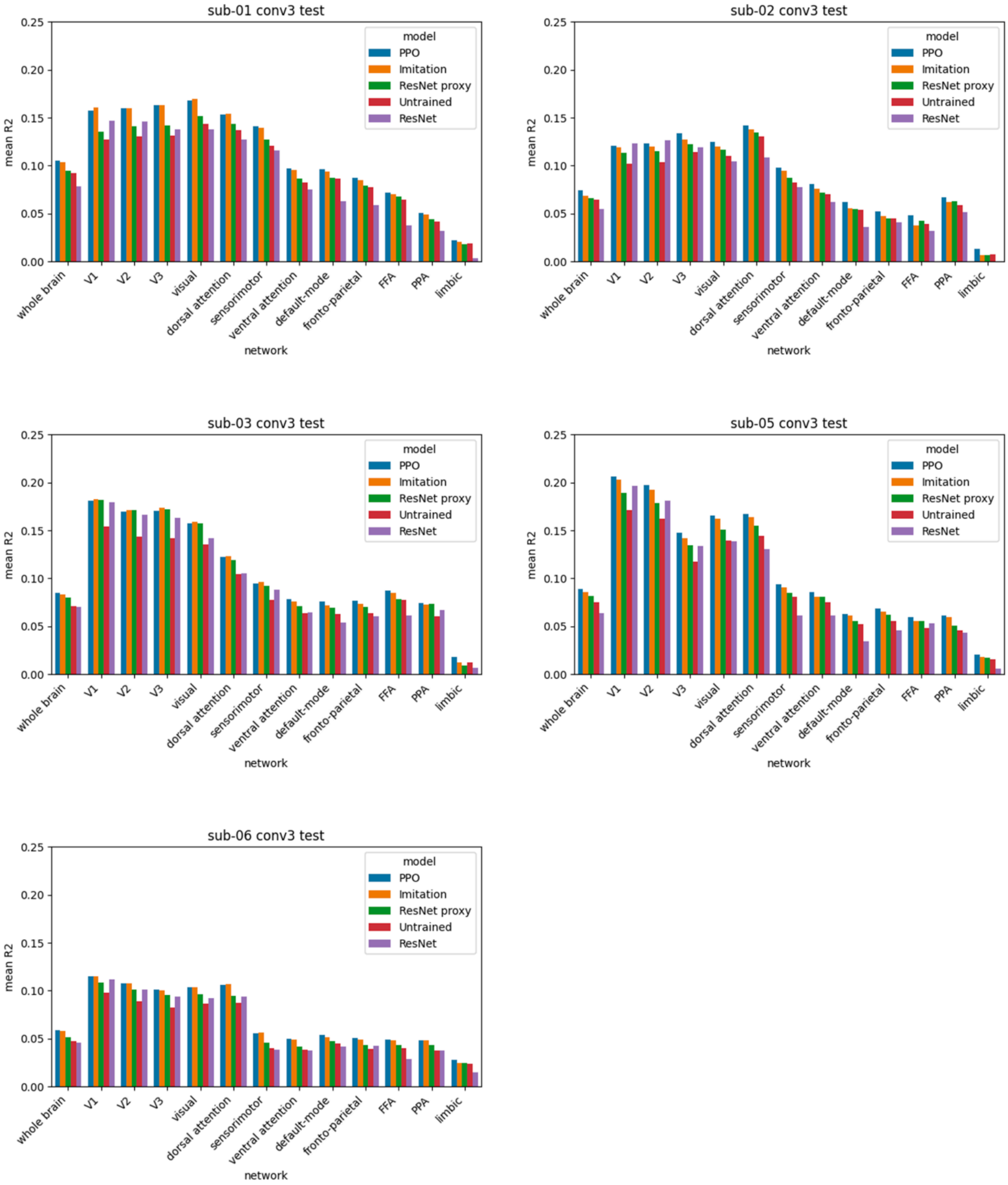
